# Relationships between depressive symptoms and brain responses during emotional movie viewing emerge in adolescence

**DOI:** 10.1101/542720

**Authors:** David C. Gruskin, Monica D. Rosenberg, Avram J. Holmes

**Affiliations:** Department of Psychology, Yale University, New Haven, Connecticut 06520, USA; Department of Psychiatry, Yale University, New Haven, Connecticut 06520, USA

**Author notes:** Corresponding author. Correspondence: Avram J. Holmes, Yale University, Department of Psychology, 402 Sheffield Sterling Strathcona Hall, 1 Prospect Street, New Haven, CT 06511.

**Keywords:** Depression, Brain development, Emotion, Naturalistic fMRI, Adolescence, Inter-subject synchronization

## Abstract

Affective disorders such as major depression are common but serious illnesses characterized by altered processing of emotional information. Although the frequency and severity of depressive symptoms increase dramatically over the course of childhood and adolescence, much of our understanding of their neurobiological bases comes from work characterizing adults’ responses to static emotional information. As a consequence, relationships between depressive brain phenotypes and naturalistic emotional processing, as well as the manner in which these associations emerge over the lifespan, remain poorly understood. Here, we apply static and dynamic inter-subject correlation analyses to examine how brain function is associated with clinical and non-clinical depressive symptom severity in 112 children and adolescents (7-21 years old) who viewed an emotionally evocative clip from the film *Despicable Me* during functional MRI. Our results reveal that adolescents with greater depressive symptom severity exhibit atypical fMRI responses during movie viewing, and that this effect is stronger during less emotional moments of the movie. Furthermore, adolescents with more similar item-level depressive symptom profiles showed more similar brain responses during movie viewing. In contrast, children’s depressive symptom severity and profiles were unrelated to their brain response typicality or similarity. Together, these results indicate a developmental change in the relationships between brain function and depressive symptoms from childhood through adolescence. Our findings suggest that depressive symptoms may shape how the brain responds to complex emotional information in a dynamic manner sensitive to both developmental stage and affective context.

## INTRODUCTION

When it comes to interpreting emotional events, truth is subjective. Across individuals, variability in state and trait psychological characteristics can influence the ways in which we attend to, interpret, and make decisions about affective information. In their pathological form, biases in emotional information processing, including increased focus on negative stimuli and overly pessimistic interpretations of emotional events, reflect core features of affective illnesses such as major depressive disorder (1–3). Depression symptom prevalence, however, is not static across the lifespan. Rather, risk for illness onset increases dramatically during adolescence (4, 5), and depression is currently the largest single contributor to adolescent disability (6). Although substantial progress has been made in identifying psychological characteristics and cognitive biases in children and adolescents that increase the likelihood of future mental health problems (7), relationships between individual responses to complex emotional stimuli, symptoms of depression, and neurodevelopmental processes remain poorly understood.

Underpinning the emergence of affective symptoms and associated illness risk, human brain development is influenced by a complex series of dynamic processes ranging from shifts in profiles of gene expression to changes in environmental pressures (8, 9). The passage from childhood to adulthood, in particular, is marked by significant social and biological transitions (10, 11), including hierarchical changes in brain structure and function that underlie the gradual development of adaptive emotion reactivity and regulation (12). As one example, the staggered development of amygdala and medial prefrontal cortex (mPFC) contributes to the prominent increase in emotional difficulties that characterizes adolescence (12). This scheduled maturation of subcortical and cortical systems is reflected in a developmental switch in amygdala-mPFC functional connectivity and alterations in amygdala responses to emotionally evocative stimuli (13–15). The characterization of age-dependent changes in brain function would provide a tremendous opportunity to understand how neurodevelopment shapes both individual differences in emotion processing and the emergence of depression.

Although pronounced in frontolimbic circuitry, developmental changes with consequences for individual differences in affective behavior are evident in widely distributed patterns of brain function. In adults, such patterns reflect individual symptom profiles of patients with depression (16, 17) and other psychiatric disorders (18, 19). Much like a fingerprint, distributed spatiotemporal patterns of brain function allow for the identification of individuals within a broader population (20) and the prediction of behavioral phenotypes across cognition, personality, and emotion (21, 22). These individual-specific functional profiles emerge over the course of development, settling into a more stable, idiosyncratic pattern during adolescence (23). Delays in this process of differentiation are associated with the expression of psychiatric symptoms, including those related to depression (23). Accordingly, data-driven approaches that consider the distributed neurobiological changes associated with adolescence can complement region-or circuit-specific work to illuminate the sources of individual variation that contribute to symptom burden and risk for depression across development.

Embedded within the gradual process of neurodevelopment, brain responses are continually changing over shorter time-scales as individuals react to, and interact with, the environment. The diverse repertoire of moment-to-moment fluctuations in brain activity and functional connectivity is instrumental for higher-order cognition (24, 25). Furthermore, impairments in functional brain dynamics have been implicated in psychiatric illness (18, 26). Although task-based and task-free fMRI (functional magnetic resonance imaging) have dominated modern research on the biological basis of depression, much of the work in these domains has relied on static analytic approaches that assume stable brain responses across time (27). An emerging dynamic paradigm that combines the targeted nature of task-based imaging with the minimal demands of resting-state research is movie-watching fMRI (28, 29). In addition to providing high-quality data by reducing head motion and increasing compliance (28, 30), movie-watching fMRI facilitates the study of naturalistic affective processing. Movies, with their dynamic, feature-rich content, can elicit emotional responses similar to those experienced in real-world situations (29, 31). A powerful approach used to study these film-induced brain responses is inter-subject correlation (ISC) (32). This method describes the similarity between multiple individuals’ brain responses to a time-locked stimulus. ISC has proven useful for identifying the possible brain bases of emotion processing in healthy individuals (33–35) and group differences in adult depression (36, 37). However, relationships between functional responses to naturalistic affective stimuli and individual differences in depressive symptoms remain poorly understood, especially in developing populations.

To characterize relationships between functional brain responses to emotional information and depression symptoms, we analyzed neuroimaging and questionnaire data acquired from 112 children and adolescents by the Healthy Brain Network project (38). First, we show that synchronization of brain activity, indexed with ISC, scales with depressive symptom severity in adolescents but not children. Next, we demonstrate that the synchrony of brain responses, as well as the relationship between this synchrony and depressive symptoms, is influenced by the emotional content of the movie. Finally, we show that patterns of movie-evoked brain activity are common between adolescents, but not children, with similar item-level depressive symptom profiles. These collective results suggest that depressive symptoms influence how the brain responds to emotional information in a dynamic manner that is sensitive to both developmental stage and moment-to-moment fluctuations in affective content.

## RESULTS

To understand relationships between naturalistic emotional processing and depressive symptoms across development, we characterized children’s and adolescents’ fMRI signal responses to an emotionally evocative clip from the film *Despicable Me* (Fig. 1; see Methods for information on data acquisition and preparation). Specifically, we asked whether the “typicality” of an individual’s blood-oxygenation-level-dependent (BOLD) signal time-courses—that is, their similarity to the rest of the group—was related to that individual’s depressive symptom severity measured with self-report responses to the Moods and Feelings Questionnaire (MFQ-SR) (39). We next asked whether relationships between spatiotemporal activity patterns and depressive symptoms were influenced by the emotional content of the film, and whether individuals with more similar patterns of brain activity reported more similar symptom patterns. Of note, our participant sample includes children and adolescents with and without diagnoses of depression and other comorbid disorders (Supplementary Table 1). Here we focus on symptom severity rather than clinical diagnosis to characterize dimensional relationships between brain function and behavior.

**Figure 1:**
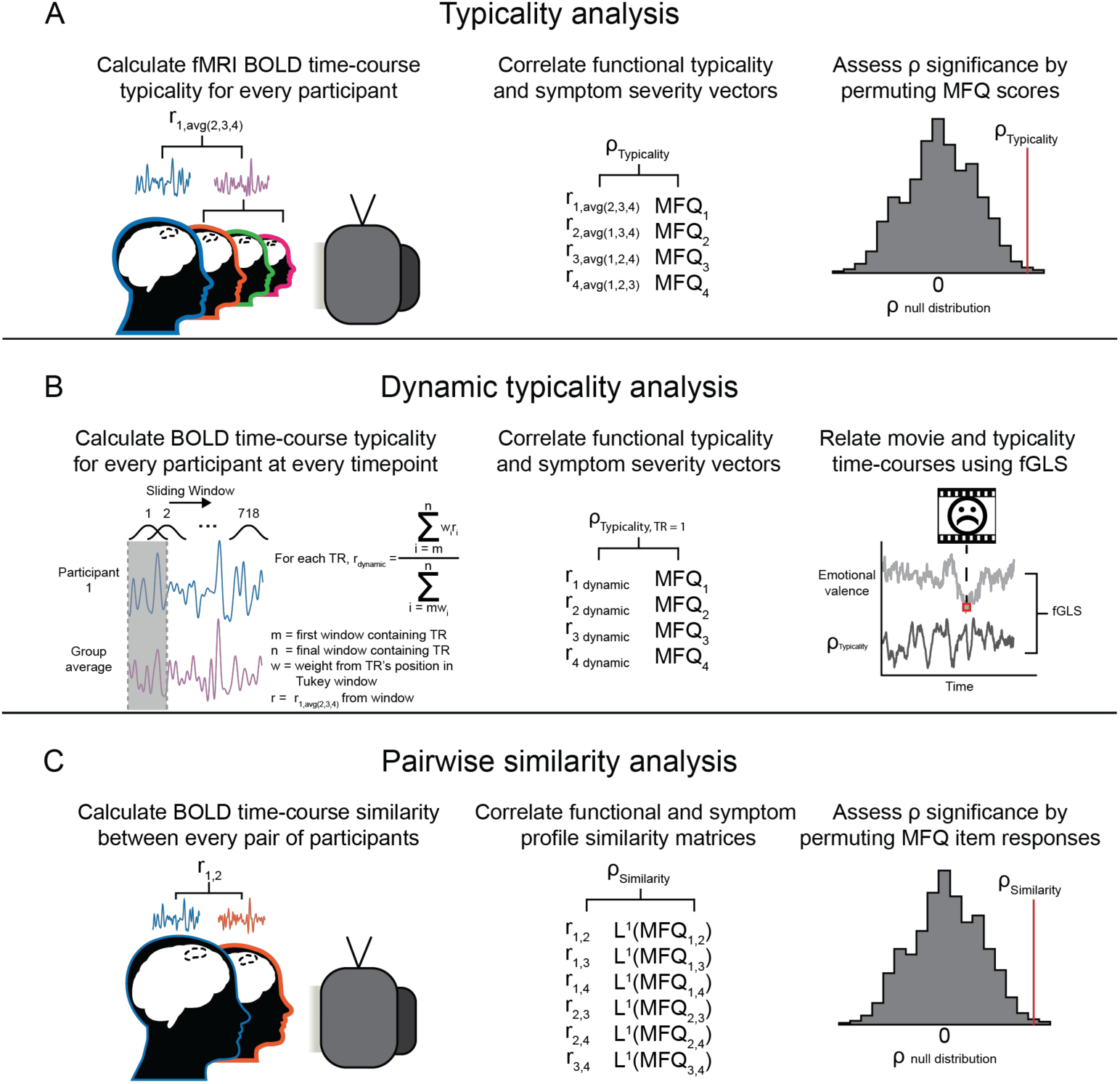
Typicality, dynamic typicality, and pairwise similarity analysis pipelines. (A) Typicality analysis: Functional typicality for each participant and each brain region (parcel) was assessed with the Pearson correlation between each individual’s BOLD response time-course and the group average time-course from the remaining participants. The resulting functional typicality vector was Spearman-correlated with depressive symptom scores. Permutation testing was used to assess significance. (B) Dynamic typicality analysis: ISC was estimated at each functional time point using a sliding window/weighted average approach (see Methods). The typicality correlation described in (A) was applied at each time point. The association between fluctuations in emotional content and the functional typicality/symptom severity relationship was evaluated using feasible Generalized Least Squares (fGLS) regression. (C) Pairwise similarity analysis: BOLD time-course similarity between every pair of participants was calculated via Pearson correlation. The resulting values were related to similarity in depressive symptom profiles through Spearman correlations and significance was determined through permutation testing.

### Greater depressive symptom severity is associated with reduced functional typicality during emotional movie viewing in adolescence

The free viewing of an emotionally evocative *Despicable Me* clip elicited stereotyped BOLD responses across participants, as measured by inter-subject correlation (Fig. 2A). Consistent with recent work from Guo and colleagues in adults (36), greater self-reported depressive symptom severity, measured with the MFQ-SR, was associated with participant-specific brain responses that were less similar to the group average (Fig. 3A). In other words, individuals with more severe depressive symptoms showed less typical fMRI response time-courses, a measure we call “functional typicality” (Fig. 1A). This inverse relationship was expressed in several regions derived from a whole-brain parcellation (40), including parcels encompassing aspects of left dorsolateral prefrontal cortex (dlPFC; ρ=−.27, p<.01, uncorrected; Fig. 3A) and bilateral medial temporal lobes (right hemisphere ρ=−.23, p<.01; left hemisphere ρ=−.28, p<.001).

**Figure 2:**
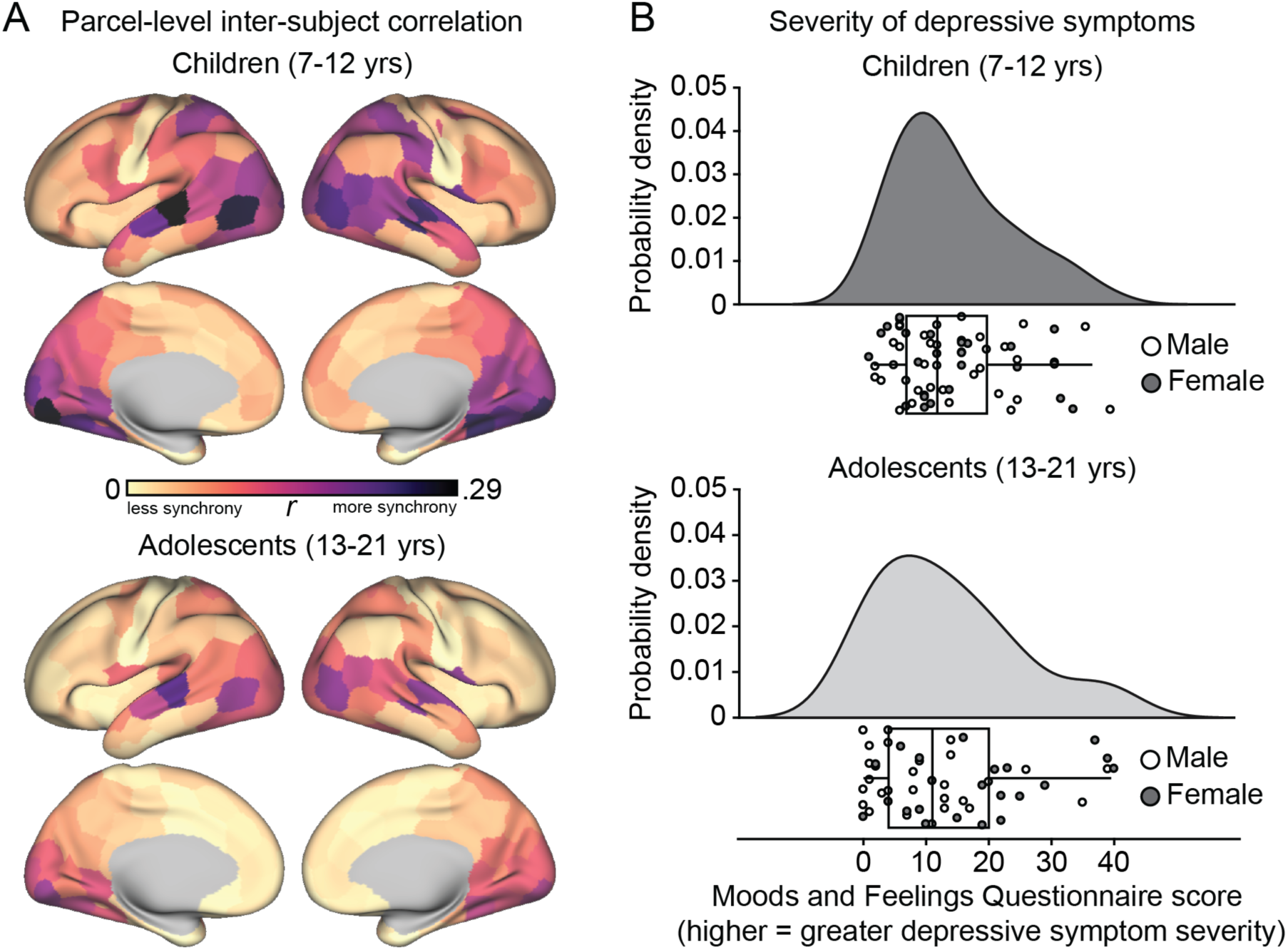
Parcel-level inter-subject correlations and depression score distributions across childhood and adolescence. (A) Inter-subject correlation analyses reveal consistent participant responses to movie viewing with a non-uniform spatial distribution across cortex. (B) Depression (MFQ-SR) score means and distributions were similar across the child (14.7±9.4 [SD]) and adolescent (13.4±11.4) groups (*t*_110_=0.65, p=0.52; Kolmogorov-Smirnov test; *D*=.20, p=.18).

**Figure 3:**
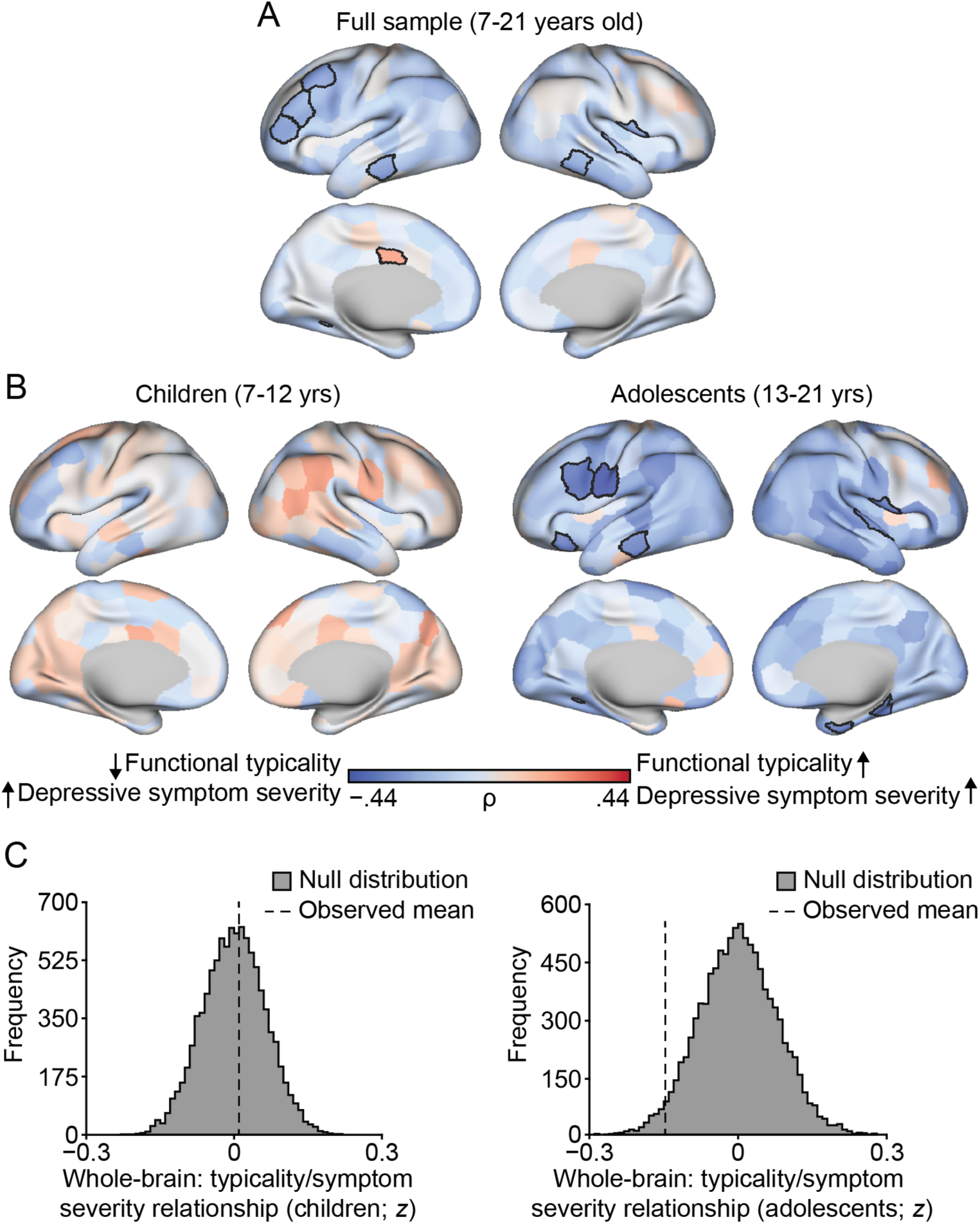
Functional typicality is related to depressive symptom severity in adolescence. (A) Spearman partial correlations controlling for age and sex reveal parcels in which less typical BOLD time-courses are associated with greater (blue) depressive symptom severity in the whole sample (bordered parcels significant at p<.01 uncorrected). (B) The same analysis shown in (A), repeated for the child and adolescent groups. (C) The histograms reflect permutation tests demonstrating that the observed mean functional typicality/symptom severity correlation coefficient across all parcels was stronger than that from the corresponding null distribution in the adolescent, but not child, group (whole-brain averages: ρ_children_=.01, p=.44; ρ_adolescents_=−.15, p<.05).

The transition from childhood to adolescence is marked by changes in brain and behavioral responses to emotionally evocative and naturalistic stimuli (14, 41, 42), including age-dependent shifts in functional synchronization. Accordingly, analyses that group children and adolescents together could fail to identify age-dependent associations. To examine how the relationship between depressive symptoms and naturalistic emotion processing changes with development, we divided the full sample into two age-groups (children: 7-12 years old, n=62; adolescents: 13-21 years old, n=50) using a split point informed by prior work (43) and repeated the functional typicality analysis separately in each group (Fig. 3B). Findings in the adolescent group mirrored those observed across the full sample, such that depressive symptom severity was negatively associated with the typicality of movie-evoked functional time-courses in a number of parcels (e.g. left orbitofrontal cortex, ρ=−.34, p<.01, left medial temporal lobe, ρ=−.34, p<.01). This relationship was consistent across the brain such that the average correlation coefficient across all parcels was significantly stronger than that of a corresponding null distribution (whole-brain average ρ=−.15, p<.05; Fig. 3C). We elected to refrain from using multiple comparison corrections as the presence of a whole-brain effect provided additional evidence for our results not captured by frequentist statistics; however, we note that in our parcel-level follow-up analyses, ∼2 false positive relationships can be expected per sub-figure (each visualizing 201 parcels) given our p<.01 threshold.

In the child group, on the other hand, no parcel-level correlations between symptom severity and functional typicality were significant at p<.01, and the mean correlation coefficient across all parcels was nominally positive (parcel-level |ρs_children_|<.29, ps>.01; whole-brain average ρ=.01, p=.44). The whole-brain relationship between symptom severity and functional typicality nominally differed between the child and adolescent groups (whole-brain average ρ_children_-ρ_adolescents_=.16, p=.06; Supplemental Figure 1). Importantly, MFQ score means and distributions as well as responses to 32 of the 33 individual scale items did not significantly differ between the child (14.7±9.4[SD]) and adolescent (13.4±11.4) groups (mean MFQ scores: *t*_110_=0.65, p=.52; Kolmogorov-Smirnov *D*=.20, p=.18; individual question *|t*-statistics*|<*1.98, ps>.05; MFQ item 13: “I was talking more slowly than usual”, *t*_110_ =2.16, p=.03), suggesting that differences in the functional typicality/symptom severity relationship cannot be explained solely by disparities in symptom severity or profiles between the two groups (Fig. 2B). Data tables detailing our results for all cortical and subcortical parcels that passed the p<.01 threshold are available in the Supplementary Materials.

### Emotional movie content increases functional synchrony

Our analyses revealed a relatively consistent pattern of brain activity across participants during movie viewing, replicating prior work in this domain (32). However, the specific stimulus features that evoke this inter-subject consistency have yet to be characterized in developmental populations. To test whether dynamic emotional movie content drives functional synchronization, we first estimated fluctuations in ISC using a Tukey sliding window approach (Fig. 1B; 24-second window, 30 time points; see Methods for details). This resulted in functional typicality scores for every participant and every fMRI volume, which we averaged across individuals to generate a dynamic ISC time-course for each parcel in our whole-brain atlas. Next, we related these dynamic ISC time-courses to independent ratings of the emotional content of the movie using feasible Generalized Least Squares (fGLS) regression, a technique sensitive to the autocorrelation present in the movie and imaging data (44).

Consistent with prior work in adults (34), emotional intensity (the absolute value of the emotional valence ratings) was positively associated with ISC in the full participant sample (age 7–21) such that brain synchrony across individuals was greater during more emotionally evocative moments of the movie (Fig. 4). This relationship was strongest in parcels encompassing aspects of frontolimbic circuitry implicated in emotion processing, including bilateral anterior insula (right hemisphere β=0.11, p<.0001, left hemisphere β=0.06, p<.01) and right anterior cingulate cortex (β=0.07, p<.01), as well as in areas implicated in cognitive control, such as right dorsolateral prefrontal cortex (β=0.09, p<.0001). Conversely, emotional valence was *negatively* associated with ISC in several parcels, including bilateral dlPFC (right hemisphere β=−0.12, p<.001; left hemisphere β=−0.07, p<.01) and anterior insula (right hemisphere β=−0.09, p<.01; left hemisphere β=−0.09, p<.01).

**Figure 4:**
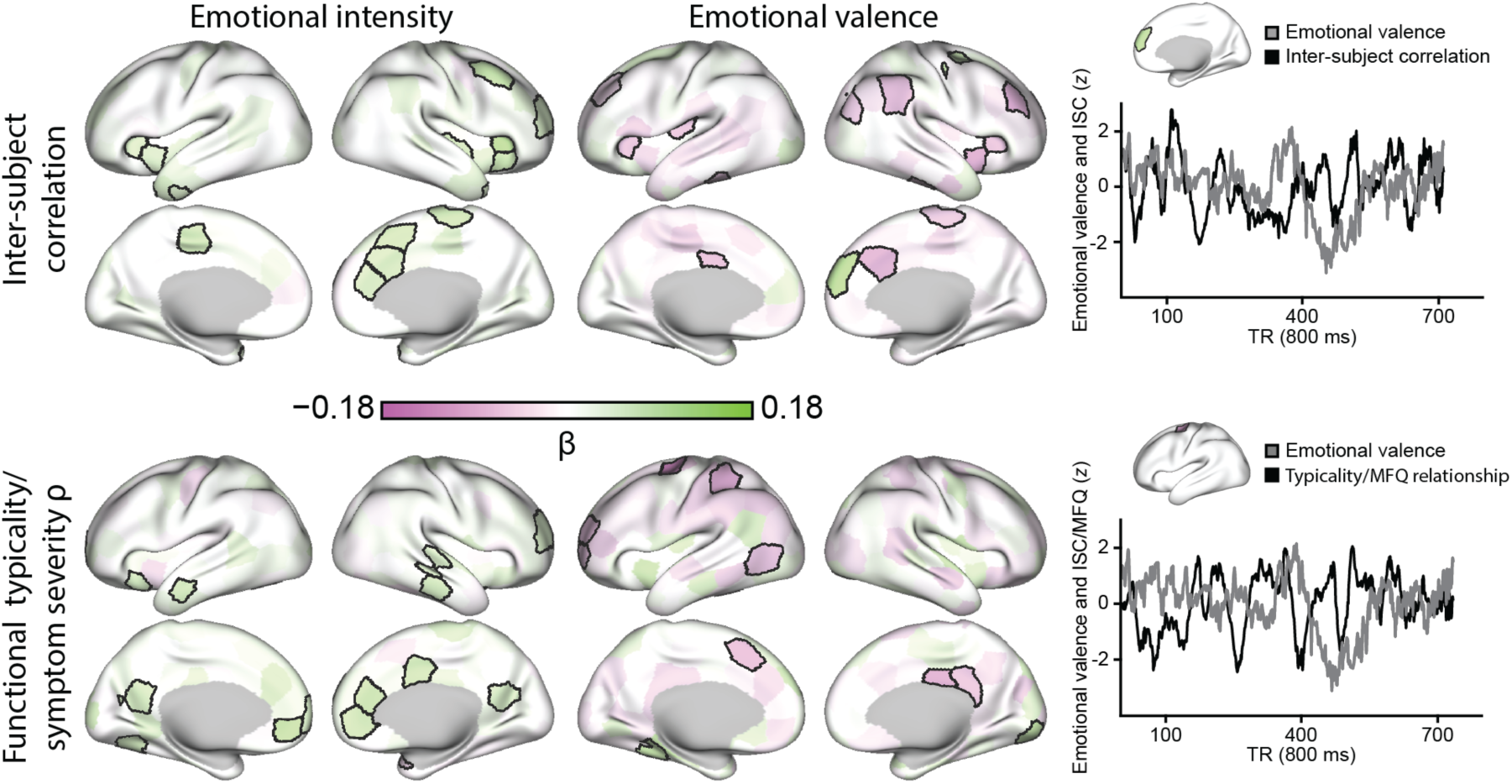
Fluctuations in emotional movie content influence both inter-subject correlations and the relationship between functional typicality and depressive symptom severity (full sample). Across the full sample, more emotionally charged moments of the movie were associated with stronger inter-subject correlations (top left) and weaker functional typicality/depressive symptom severity relationships (bottom left), as determined by fGLS regression (parcel threshold p<.01). BOLD synchronization was higher in several parcels when the clip was more negatively valenced (top middle). Positively valenced moments of the movie maximized the inverse relationship between functional typicality and symptom severity in more parcels than did negatively valenced moments (bottom middle). Line plots showing the ISC and functional typicality fGLS regressions are included for visualization purposes (right).

When independently considering children and adolescents, the dynamic ISC findings were similar across both groups, with positive inter-group spatial correlations for both ISC/emotional intensity (ρ=.20, p<.001) and ISC/emotional valence relationships (ρ=.16, p<.01; see Supplementary Figure 2 for age-group-specific maps). While the topographic distributions of effects were preserved across the two age-groups, differences in effect size were apparent (Supplementary Figure 3) such that the relationship between increased synchronization and more negatively valenced content was stronger in adolescents than in children. This age-related difference was notable in several regions including bilateral mPFC (right hemisphere *z*=3.33, p<.001; left hemisphere *z*=3.33, p<.001) and right temporoparietal junction (*z*=2.9, p<.01). These results suggest that emotional film content synchronized brain responses in both an age-invariant (topography) and an age-specific (effect size) manner.

### Emotional movie content weakens the relationship between ISC and depressive symptom severity in adolescence

Developmental studies have reported information processing biases in depression, including increased engagement with negative information and excessively negative interpretations of emotional events (45, 46). Our analyses suggest that inter-subject synchronization is related to the emotional content of the stimulus. Given that individuals with depression can exhibit emotion-driven attentional and cognitive biases, the atypical BOLD time-courses we observed in more depressive individuals may reflect dynamic biases in attention to, appraisal of, or difficulty disengaging from emotional information. Consequently, we hypothesized that the emotional movie content would dynamically influence the relationship between functional typicality and depressive symptom severity.

Using the dynamic ISC time-courses described above (Fig. 1B), we first calculated the relationship between functional typicality and symptom severity at each time point. We next related the resulting time-courses of correlation coefficients (reflecting the fluctuating strength of the functional typicality/symptom severity relationship) to the emotional intensity and emotional valence time-courses using fGLS. The functional typicality/symptom severity relationship was stronger (more negative) during less emotional moments of the movie in parcels encompassing aspects of bilateral vmPFC (right hemisphere β=0.08, p<.001, left hemisphere β=0.09, p<.01; Fig. 4) and posterior cingulate cortex (right hemisphere β=0.06, p<.01, left hemisphere β=0.07, p<.01). Similar to the dynamic ISC results, emotional valence was negatively associated with the strength of the functional typicality/symptom severity relationship in several parcels, including left dmPFC (β=−0.09, p<.01) and right posterior cingulate cortex (β=−0.09, p<.01).

### Adolescents with similar depressive symptom profiles share more similar brain responses during emotional movie viewing

Our results demonstrate that functional typicality during movie watching is related to depressive symptom severity in adolescence. However, depression is a heterogeneous syndrome and the assessment of gross depressive symptom severity can mask the presence of diverse symptom profiles. This is especially the case during development, when substantial variability in emotional, social, and cognitive maturity is thought to give rise to diverse clinical presentations (47). Recent work suggests that different depressive symptom profiles track with distinct patterns of brain function (16, 17) and cognitive impairments (48). Accordingly, two individuals who share similar levels of depressive symptom severity might express qualitatively different symptom profiles and associated brain phenotypes.

To determine whether BOLD time-courses during emotional movie watching index depressive symptom profiles, we used the L1 (Manhattan) distance to investigate whether pairs of individuals who were more alike in their item-level responses to the MFQ-SR also exhibited more similar functional time-courses (Fig. 1C). In the full sample (age 7–21), this pairwise similarity analysis revealed that participant pairs with more similar symptom profiles were also more similar in terms of their movie-induced BOLD activity in a subset of parcels (controlling for similarity in age and sex; Fig. 5A). The hypothesized global effect was present in the adolescent group (whole-brain average ρ=−.06, p<.05; Fig. 5C), where it was preferentially expressed in bilateral anterior insula (right hemisphere ρ=−.08, p<.01, left hemisphere ρ=−.09, p<.01; Fig. 5B), bilateral orbitofrontal cortex (right hemisphere ρ=−.10, p<.01, left hemisphere ρ=−.10, p<.01), and right amygdala (ρ=−.09, p<.01), as well as in right hippocampus (ρ=−.19, p<.01). Analogous to our typicality findings, there were no global or parcel-level relationships between symptom profile and functional similarity in children (whole-brain average ρ_children_=.004, p=.46; parcel-level |ρs_children_|<.16, ps>.01), resulting in a nominal difference in whole-brain effects between the two groups (whole-brain average ρ_children_-ρ_adolescents_=.06, p=.06; Supplemental Figure 1). Of note, the relationships between symptom profile similarity and functional time-course correspondence in adolescents were to some degree accounted for by similarity in symptom severity, operationalized as the absolute value of the difference between two participants’ MFQ scores (whole-brain average of symptom profile/time-course similarity relationship controlling for symptom severity similarity ρ=−.03, p=.12). However, time-course similarity was nominally more related to similarity in symptom profiles than symptom severity (ρ=−.06 vs ρ=−.05), suggesting that while depressive symptom severity and profile similarity are largely overlapping, symptom profiles may reflect additional unique aspects of brain function.

**Figure 5:**
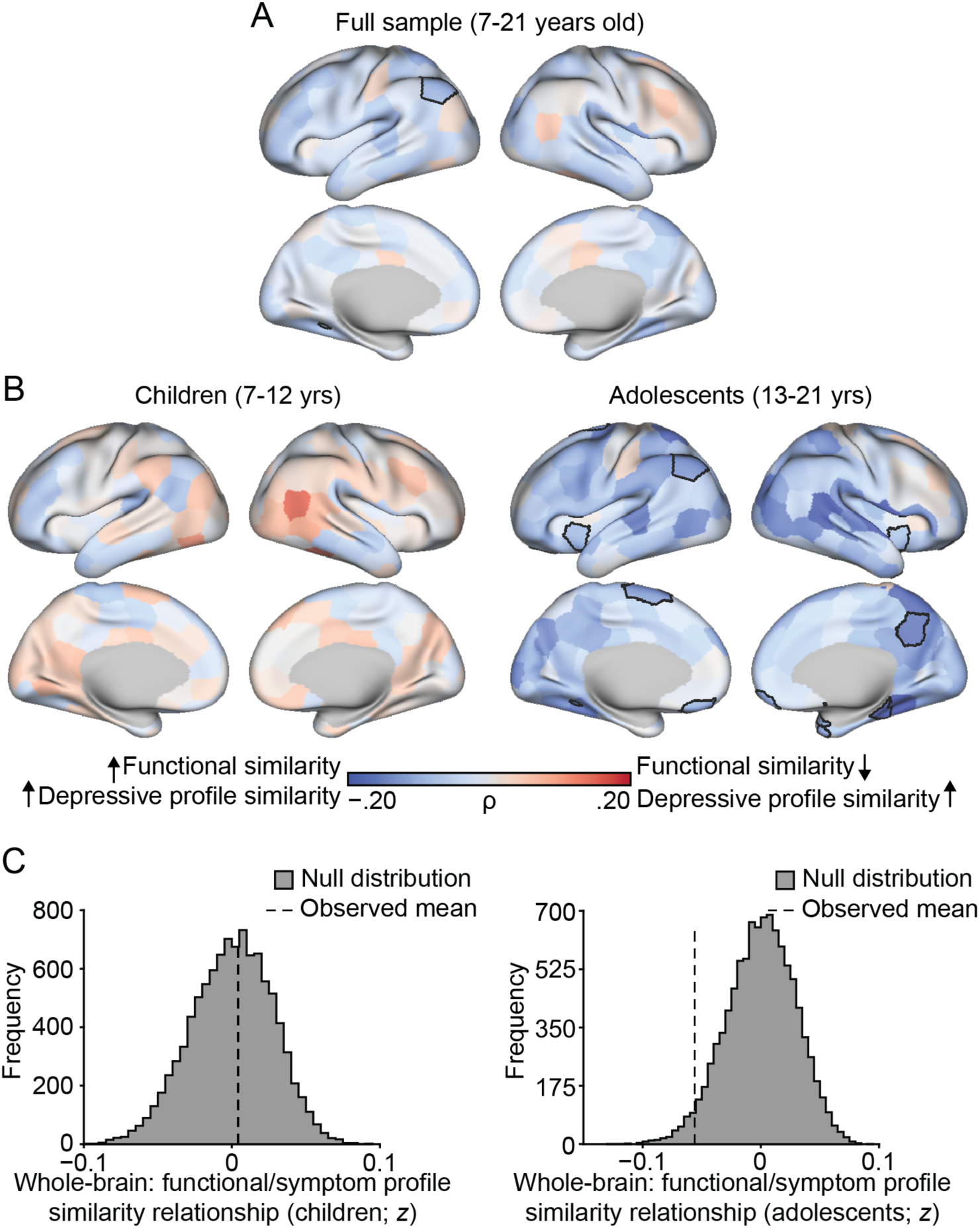
BOLD time-course similarity scales with depressive symptom similarity in adolescents. Spearman partial correlations controlling for similarity in age and sex reveal parcels in which pairwise BOLD time-course similarity is associated with more similar (blue) depressive symptom profiles in the whole sample (bordered parcels significant at p<.01 uncorrected). (B) The same analysis shown in (A), repeated for the child and adolescent groups. (C) The histograms reflect permutation tests demonstrating that the observed mean functional similarity/symptom profile similarity correlation coefficient across all parcels was stronger than that from a corresponding null distribution in the adolescent, but not child, group (whole-brain averages: ρ_children_=.004, p=.46; ρ_adolescents_=−.06, p<.05).

## DISCUSSION

Children and adolescents with severe depressive symptoms often describe viewing the world around them through a filter, focusing on negative information and forming overly pessimistic interpretations of the thoughts and actions of others (46). A fundamental question facing human neuroscience is how these emotional information processing biases and the associated expression of depression symptoms emerge during development. Here we extend research on the brain mechanisms underlying this phenomenon. First, we demonstrated that greater depressive symptomatology was associated with less typical brain responses to an emotional clip from the film *Despicable Me* in adolescents but not children. Second, we established that the strength of this brain/behavior relationship scaled with the moment-to-moment affective content of the movie, such that the depressive symptom severity and functional typicality association was more prominent when the movie was less emotional. Finally, we found that adolescents with more similar depressive symptom profiles shared more similar brain responses to the movie. Collectively, these results highlight time-varying features of brain function and emotional information processing that reflect the presence and severity of depression symptoms across development.

The transition from childhood to adulthood is characterized by changes in brain function and structure that constrain moment-to-moment responses to affective information (10, 12, 49). In both children and adolescents, depression is associated with altered brain responses to static emotional images (1, 14, 50). In adults, patients with melancholic depression exhibit reduced group-level inter-subject functional synchronization during film viewing (36). Building upon this literature, our ISC typicality and similarity analyses revealed that adolescents with more severe depressive symptoms exhibited less typical brain-wide responses during emotional movie viewing, and that these responses were more similar in pairs of individuals who were more alike in their symptom profiles. These effects were particularly evident in several regions commonly implicated in affective integration and regulation (e.g. orbitofrontal cortex, anterior insula, amygdala) (51). Prior work suggests that specific changes in amygdala– mPFC functional connectivity during emotional face viewing underlie shifts in psychiatric symptoms during the transition to adulthood (13). The distributed relationships between patterns of brain function and depression observed here provides novel insight into the neurobiological basis of affective processing in adolescents and suggests an intriguing model for the more general development of emotion-relevant regulatory systems throughout the brain.

Although much of the current literature treats emotion processing as time-invariant by focusing on average responses to static stimuli, the brain possesses a dynamic organizational structure that adjusts in response to explicit task demands. Here, we establish the importance of dynamic approaches to the study of emotion processing by demonstrating that the observed relationship between functional typicality and depressive symptom severity was sensitive to the emotional content of the movie. More specifically, the inverse functional typicality/symptom severity relationship was weaker when the movie was more emotionally charged. This effect was noted in in several higher-order brain regions, including bilateral vmPFC and PCC. Recent studies have found that functional similarity in vmPFC and PCC/posteromedial cortex increases when individuals share similar interpretations of an event (52, 53). Based on this literature, participants in our study with greater depressive symptoms may have formed more similar interpretations of the movie when it was more emotional, leading to relative increases in ISC, thereby weakening the inverse functional typicality/symptom severity relationship. Similarly, these participants may have engaged in more stimulus-independent cognition and/or idiosyncratic interpretation during less emotional (and perhaps less engaging) moments, which could account for the inverse association between functional typicality and symptom severity observed across the clip. Although speculative, this candidate explanation aligns with interpretation bias, a phenomenon in which depressed individuals adopt rigid, overly pessimistic responses to emotional (and especially negatively valenced) information (3, 46, 54, 55). This set of findings deserves close consideration in future studies, as interpretation bias represents just one possible explanation for these observed effects. For example, it may be the case that brain function and symptom severity were less related during emotionally negative compared to positive moments of the clip because more depressed individuals may preferentially attend to negative stimuli, regardless of how those stimuli are interpreted.

Psychiatric research has increasingly focused on a dimensional perspective of illness that incorporates transdiagnostic conceptions of neurobiology and behavior (56). This approach is particularly relevant to the study of major depressive disorder, a heterogeneous syndrome with hundreds of possible clinical presentations (57). Our finding that functional similarity in adolescents was nominally more associated with depressive symptom profiles than symptom severity suggests that an individual’s brain responses to naturalistic stimuli may be sensitive to their specific pattern of symptoms. Understanding the neurobiological signatures of depressive symptom profiles in children and adolescents is especially crucial, as both the prevalence and consequences of depression are magnified during the transition to adulthood (46). Here, the presence of relationships between time-course typicality/similarity and depressive symptoms in adolescents and not children may reflect effects of symptom duration and/or developmental stage. More specifically, it is possible that inter-subject synchrony scales with symptomatology once individuals have spent a considerable amount of time living with depressive symptoms, irrespective of age. It is impossible to confirm or counter this explanation without being able to control for clinical features such as age of onset and time since last episode. However, the recent discovery of similar adolescent-emergent relationships between neurogenetic/functional profiles and psychiatric symptomatology (23, 58) suggests that our present findings could be related to developmental stage. One such relationship was identified by Kaufmann and colleagues, who reported that while adolescents with increased overall psychiatric symptom severity exhibited decreased resting-state connectome distinctiveness compared to healthy individuals, no significant relationship between these variables was found in children (23). Although the association between functional brain typicality and symptom severity was negative in our analyses, we note that functional typicality has been shown to scale positively with some clinically relevant features (trait paranoia (59)) and negatively with others (depression (36) and autism (60) symptom severity). Therefore, while functional typicality appears to serve as an index of psychopathology that emerges during adolescence, future work should clarify how these relationships may differ across psychiatric diagnoses and analytic approaches.

Several limitations should be considered when evaluating the current findings. First, individual differences in ISC vary significantly with the specific characteristics of the stimulus and population being studied (29, 30). This is particularly relevant to the study of emotional processing, as films with different affective structures might reveal distinct relationships between depressive symptoms and brain function. Ongoing deep-phenotyping efforts involving a diverse array of stimuli and behavioral assays may be helpful in illuminating general relationships between affective brain response and behavior (61). Second, the emotional content of the movie was rated by six independent adults (mean age=25.5 years old) whose affective experiences while viewing the movie might have qualitatively differed from those of the participants. Although this would likely only hinder our ability to relate emotional film content to brain function, it is difficult to predict exactly how this might influence our analyses. Finally, the design of this study prevented us from identifying specific cognitive and affective processes associated with the ISC/depressive symptom relationships and how these relationships might change within individuals across development. These findings motivate longitudinal research to examine how age-related changes in individuals’ brain responses to naturalistic emotional stimuli relate to depressive symptoms across development. Such efforts hold considerable promise in advancing both our ability to predict clinical outcomes and our understanding of neurobiological mechanisms underlying psychiatric illness (22, 62).

## CONCLUSIONS

How the presence and severity of psychiatric symptoms colors emotional experiences across development is a central and challenging question in affective neuroscience. Our data suggest that atypical brain responses to an emotional movie may constitute functional markers of depression that emerge in adolescence and serve as signatures of item-level depressive symptom profiles. The sensitivity of the observed functional typicality/symptom severity relationship to the dynamic affective content of the movie is consistent with the proposed core role of affective information-processing biases in depression (3). Furthermore, the preferential expression of these effects in frontolimbic areas aligns with an extensive literature implicating these regions in the processing of static emotional images (1, 51, 63). However, the presence of whole-brain relationships suggests that naturalistic stimuli may afford the opportunity to characterize qualitatively different patterns of activity in response to emotional content. Due to the method’s ecological validity (29), ability to provide high quality data (28, 30), and support of sophisticated analytic techniques (28), we join a growing number of researchers in recommending that naturalistic paradigms be used to study the brain bases of psychiatric disorders. To conclude, our discovery of a developmental change in the relationships linking brain function and depressive symptoms encourages further development of naturalistic and biologically informed methods for the early detection and prevention of psychiatric illness across development.

## METHODS

Neuroimaging and questionnaire data from 563 participants were downloaded from the data portal for the Child Mind Institute’s Healthy Brain Network (HBN) project (38), a large, ongoing initiative that has been collecting data from a high-risk community sample of children and adolescents with perceived clinical concern. All participants provided written consent or assent, and consent was obtained from the parents or legal guardians of participants younger than 18 years old. The HBN project was approved by the Chesapeake Institutional Review Board. Of all the participants whose anatomical scans passed visual inspection (n=324) and had complete *Despicable Me* data (n=313), those with acceptable levels of head motion (defined a priori as maximum head displacement <3 mm or mean frame-to-frame head displacement <.15 mm on the functional scan; n=153) were included in our analyses. Finally, participants without self-report scores on the Moods and Feelings Questionnaire (n=41) were excluded, leaving 112 participants (65 F; mean age=12.7±3.3 years, range=7-21 years) for the final analyses.

### MRI data collection

All MRI data used in this study were collected at the HBN Rutgers University Brain Imaging Center site on a Siemens 3T Tim Trio magnet. Key parameters for the functional scan are as follows: TR=800ms, TE=30ms, # slices=60, flip angle=31°, # volumes=750, voxel size=2.4mm. Complete information regarding the scan parameters used for the Healthy Brain Network project can be found at: http://fcon_1000.projects.nitrc.org/indi/cmi_healthy_brain_network/mri_protocol.html#mri-scan-parameters.

### Depressive symptom inventory

Self-report responses to the Moods and Feelings Questionnaire (MFQ-SR) were used to quantify depressive symptom severity and profiles. The MFQ-SR is a 33-item questionnaire used to inventory core depressive symptomatology in clinical and sub-clinical pediatric populations (39). All HBN participants over eight years of age as well as a subset of seven-year-olds were administered the MFQ-SR. Missing item-level responses (3/3696) were imputed based on the mean response across the full sample. Mean MFQ-SR scores did not differ between the child and adolescent age-groups (child: M=14.7±9.4; adolescent: M=13.4±11.4; *t*_110_=0.65, p=0.52), and a Kolmogorov-Smirnov test revealed no significant difference in score distributions between the groups (D=.20, p=.18). Independent samples t-tests revealed that mean item-level responses were indistinguishable between groups on all but one question (individual question |*t*-statistics|<1.98, ps>.05; MFQ item 13 [“I was talking more slowly than usual”]: *t*_110_ =2.16, p=.03), suggesting that average depressive symptom profiles were consistent across the two groups. Although the MFQ-SR authors do not recommend a specific diagnostic cut-off, previous work has determined that a score of 29 can distinguish individuals experiencing a current major depressive event (64).

All participants were also administered the KSADS-COMP (65), a semi-structured DSM-5-based psychiatric interview, by a licensed clinician. Consensus DSM-5 diagnoses were then generated for each participant following the completion of the clinical interview and other study procedures. In the adolescent group, 12/50 of the participants received at least one depressive disorder diagnosis, as opposed to 6/62 of the participants in the child group. Because both groups shared similar depression score distributions but differed in their proportions of diagnoses, we calculated Cronbach’s alpha for MFQ-SR responses in both groups as a way of ensuring that our participants were accurate self-reporters of their symptoms. Internal reliability was high for both groups (child: alpha =.86, adolescent: alpha =.94). Data reflecting symptom onset were unavailable at the time of writing, but disease duration data were available for participants who entered the HBN with a depression diagnosis and history of treatment (6/112). Of these six participants, five were in the adolescent group (mean disease duration=2.46 years) and one was in the child group (disease duration=4.5 years).

### Movie clip and emotional content ratings

Functional MRI data were collected while participants viewed a ten-minute clip from the movie *Despicable Me* (01:02:09–01:12:09; presentation details available at http://fcon_1000.projects.nitrc.org/indi/cmi_healthy_brain_network/mri_protocol.html#mri-movies) (28). Six adult raters provided continuous ratings of the emotional valence of the clip on a scale from 1 to 9 (extremely negative to extremely positive) three times each using a custom PsychoPy (v1.9.0) script (66). The 18 rating runs were individually *z*-scored and averaged across raters and within TRs to yield an emotional valence vector. An emotional intensity vector was then generated using the absolute value of the emotional valence vector. To control for the effects of non-emotional aspects of the film on ISC, one rater (author D.G.) recorded the number of faces present on the screen at each TR using a PsychoPy script (averaged across three viewings). Visual and auditory intensity (brightness and loudness) of the movie were also calculated at each TR using the SaliencyToolbox (67) and a custom Matlab script (which operationalized volume as dynamic peak signal amplitude of the clip’s audio), respectively. Finally, the emotion, face, brightness, and loudness time-courses were shifted by 6 TRs (4.8s) to account for hemodynamic lag.

### fMRI data preprocessing

Functional data were preprocessed in AFNI (68). Three volumes were first removed from the start of each ten-minute run. Functional images were then despiked, corrected for head motion, aligned to the corresponding skull-stripped anatomical image with a linear transformation and then to the MNI atlas via nonlinear warping, and spatially smoothed with a 4 mm full-width at half-maximum filter. Covariates of no interest, including a 24-parameter head motion model (6 motion parameters, 6 temporal derivatives, and their squares) and mean signal from subject-specific eroded white matter, ventricle, and whole-brain masks were regressed from the data. Data were band-pass filtered from .01 to .1 Hz. Voxel-wise BOLD signal time-courses were averaged within regions of interest using a 268-node whole-brain parcellation in order to maximize power for a pairwise similarity analysis. MNI coordinates and network affiliations for all of the parcels mentioned in this study can be found at https://bioimagesuiteweb.github.io/webapp/connviewer.html.

### Static inter-subject correlation (ISC) analyses

### Functional typicality

For every parcel, each participant’s BOLD signal time-course was *z*-standardized and Pearson correlated with the average BOLD signal time-course from the rest of the group (Fig. 1A). This procedure resulted in a 112 subject x 268 parcel typicality matrix. The columns of this typicality matrix were related to MFQ-SR scores through Spearman partial correlations (controlling for age and sex) to reveal parcels in which BOLD signal time-course typicality was associated with depressive symptom severity.

Given the dependency structure of the correlation coefficients generated by ISC analyses (69), statistical significance of the relationships between BOLD signal typicality and depressive symptom severity was evaluated using nonparametric significance testing. First, the order of the questionnaire variables was randomly shuffled such that one participant’s typicality score was paired with a random participant’s MFQ-SR score, age, and sex. Spearman partial correlations were then performed and the process was repeated 10,000 times to generate a null distribution, and the reported p values were calculated using the following one-tailed formula (listed below for a positive Fisher’s *z*-value):

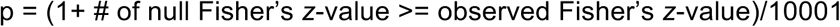

In order to test the extent to which the relationship between functional typicality and depression score was consistent across the whole brain, we averaged the Fisher’s *z*-values obtained for each parcel in the real data to get the observed mean rho value. We then computed the same average *z*-value for each permutation of the null data and performed the following one-tailed test (listed below for a positive Fisher’s *z*-value):

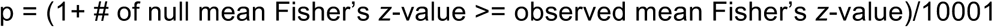

### Pairwise symptom profile similarity

For every possible pair of 112 participants, a phenotypic matrix was constructed by finding the L1 (Manhattan) distance, a measure appropriate for ordinal and trinary data (e.g. Chekroud et al., 2017 (70); high L1 distance=less similar), between each pair’s item-level responses to the MFQ (Fig. 1C). Next, parcel-level BOLD time-courses from the first participant were *z*-standardized and Pearson correlated with corresponding time-courses from the second to create a 112 x 112 BOLD similarity matrix for every parcel. Correlation coefficients were Fisher z-transformed, averaged across all subject pairs, and transformed back to *r* values to generate the average ISC maps shown in Fig. 2A. Spearman partial correlations (controlling for differences in age and sex) were then used to reveal cortical parcels in which more similar BOLD time-courses are associated with more similar depressive symptom profiles. Significance for these results was tested using a permutation design analogous to the one used in the typicality analyses. Specifically, each participant pair’s item-level MFQ responses, as well as their similarity in age (absolute value of the difference between the two participant’s ages) and sex (binary classification for same vs. different), were associated with a random pair’s functional similarity correlation coefficient, and the analysis was repeated 10,000 times.

Differences in the static ISC/depressive symptom maps (typicality and similarity; Supplemental Figure 1) for the two age groups were assessed by first Fisher transforming the observed correlation coefficients and the null coefficients generated during permutation testing. For each parcel, the adolescent group’s Fisher’s *z*-value was subtracted from that of the child group, and the significance of this difference was evaluated using the following one-tailed formula:

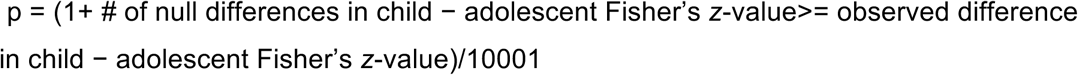

Fisher’s z-values were then transformed back to rho values for visualization.

### Dynamic inter-subject correlation analyses

#### Dynamic ISC

In order to evaluate how the content of the emotional movie drives changes in neural synchronization across participants, parcel-level dynamic ISC (i.e., functional typicality) time-courses were calculated for every participant using a tapered cosine sliding window approach (width=30 TRs, taper=15 TRs; Fig. 1B). The typicality analysis detailed above was initially performed on the first 24 seconds of the time-courses and then repeated after sliding the window by 1 TR until the end of the clip was reached. The sliding window parameters chosen for this analysis were motivated by prior research and represent a balance between capturing enough time points to calculate stable correlation coefficients and keeping the window short enough to isolate transient ISC dynamics (42). The resulting ISC time-courses were then Fisher transformed to generate a 268 parcel x 718 TR matrix for each subject reflecting their parcel-level fluctuations in synchrony with the rest of the group over the course of the movie clip.

To relate the windowed ISC time-courses to the movie vectors (emotional intensity and emotional valence, controlling for the number of faces on the screen, brightness, and loudness), estimates of ISC at each TR were obtained by first identifying every window in which a given TR was included and then taking the weighted average of those windows’ ISC *z*-values (with the weights being derived from the tapered window) (42). This approach yielded a 268 parcel x 747 TR matrix for each subject containing dynamic ISC time-courses for every parcel. The parcel x TR matrices were then *z*-standardized, averaged across participants, and zero-padded with three columns to account for the TRs discarded at the beginning of the run. The first and final 15 TRs were excluded from the following analyses as the ISC estimates for these TRs were based on a small number of windows, and 6 TRs were removed to account for hemodynamic delay, yielding 714 TRs for regression analysis.

Given the high degree of autocorrelation present in the movie feature and ISC time-courses, feasible Generalized Least Squares regression (fGLS) was used to identify how ISC scales with the emotional content of the movie. FGLS accounts for autocorrelation by minimizing the squared Mahalanobis distance of the ordinary least squares (OLS) error terms based on an estimation of their variance-covariance matrix, and GLS has previously been used to work with movie/emotion time-courses (44). A fourth-order autoregressive model was used to estimate the covariance structure of the OLS error terms based on a visual inspection of the partial sample autocorrelation functions of the OLS residual time series according to the Box-Jenkins methodology (71). Importantly, because relationships between clinical symptoms and high-level features of the stimulus such as emotional content (here, emotional intensity and valence) can be confounded by lower-level features such as visual intensity (72), time-courses of low-level stimulus features (brightness and loudness) as well as baseline social content (faces) were also included in the regression. A *t* distribution (709 degrees of freedom) was used for two-tailed significance testing of the standardized regression coefficients.

#### Dynamic typicality

The typicality/symptom severity analysis detailed above was repeated at each TR using the window-derived ISC estimates (Fig. 1B). The resulting time-courses (714 TRs) were then entered into two fGLS regression models to characterize how fluctuations in the strength of the typicality/symptom severity relationship are related to the moment-to-moment emotional intensity and valence of the movie (controlling for brightness, loudness, and baseline social content). Age-related differences in the effects of emotional movie content on ISC and the ISC/symptom severity relationship were evaluated by dividing the child – adolescent difference in standardized regression coefficients by the square root of the sum of the coefficients’ squared standard errors. Two-tailed significance testing of the resulting *z* scores was conducted against a normal distribution.

All analyses conducted in Matlab (v2016b). Data visualizations were performed using Connectome Workbench (v1.3.1) (73).

## ACKNOWLEDGMENTS

Analyses were made possible by the high-performance computing facilities provided through the Yale Center for Research Computing. We thank BJ Casey for her invaluable feedback on earlier versions of this manuscript, Kevin Anderson for assisting with visualizations, and Syntia Hadis, Erica Ho, Rowena Chin, and Meghan Collins for their help rating the emotional content of *Despicable Me*.

## Abbreviations

BOLD: blood-oxygen-level-dependent
dlPFC: dorsolateral prefrontal cortex
dmPFC: dorsomedial prefrontal cortex
fGLS: feasible Generalized Least Squares
fMRI: functional magnetic resonance imaging
ISC: inter-subject correlation
MFQ-SR: Moods and Feelings Questionnaire (Self-Report)
OLS: ordinary least-squares
PCC: posterior cingulate cortex
TR: repetition time
vmPFC: ventromedial prefrontal cortex

## AUTHOR CONTRIBUTIONS

Designed research: DCG, MDR, AJH

Curated and preprocessed data: MDR, DCG

Conducted analyses: DCG, MDR

Wrote the paper: DCG, MDR, AJ

