## Supplemental File for "Relationships between depressive symptoms and brain responses during emotional movie viewing emerge in adolescence"

*Corresponding author.

**Table S1: Participant Demographics**

|  | **Children** | | **Adolescents** | | **T-test / Chi-square** | |
| --- | --- | --- | --- | --- | --- | --- |
|  | **Mean±SD** | ***n*** | **Mean±SD** | ***n*** | **Test statistic** | ***p* value** |
| **Age** | 10.1±1.4 | 62 | 15.8±1.9 | 50 | N/A | N/A |
| **MFQ-SR** | 14.7±9.4 | 62 | 13.4±11.4 | 50 | *t*_110_ = 0.65 | .52 |
| **Sex (M:F)** | — | 25:37 | — | 22:28 | **χ**^2^_1,112_ = 0.15 | .70 |
| **Clinician: no dx** | — | 10 | — | 6 | **χ**^2^_1,112_ = 0.39 | .53 |
| **Clinician: depression dx** | — | 6 | — | 12 | **χ**^2^_1,112_ = 4.2 | <.05 |
| **Past depression dx** | — | 1 | — | 5 | **χ**^2^_1,112_= 3.84 | <.05 |
| **Current psych medication** | — | 10 | — | 12 | **χ**^2^_1,112_= 1.09 | .30 |

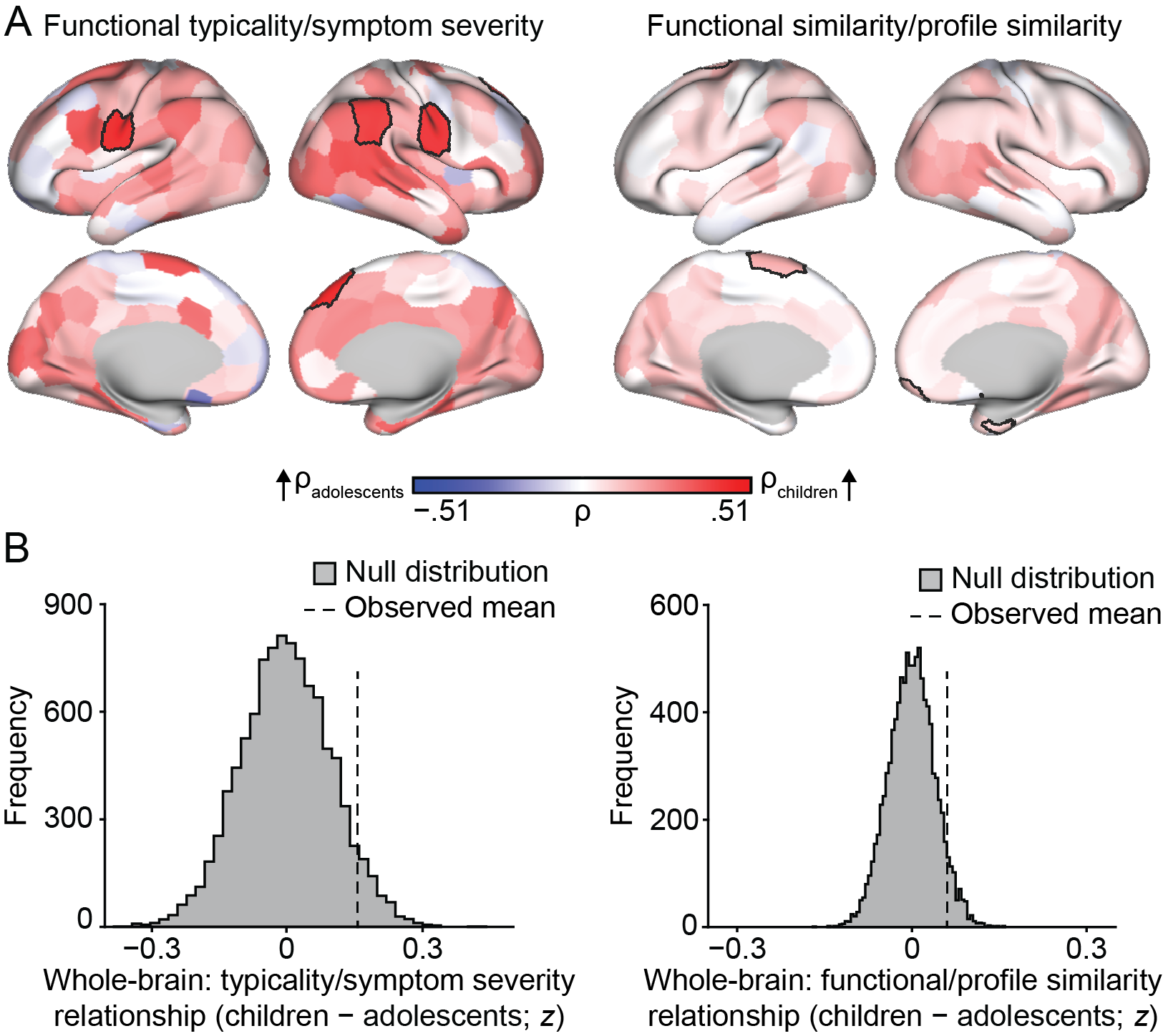

**Figure S1: Age-group differences (children – adolescents) in the ISC/depressive symptom relationships**. (A) Subtracting the ISC/depressive symptom correlation coefficients (typicality and similarity) obtained in the adolescent group from those of the child group reveals parcel-level age-group differences in the relationships visualized in Figures 3 and 5 (bordered parcels significant at p<.01). (B) The histograms reflect permutation tests showing the whole-brain age-group differences in the relationships between functional typicality/symptom severity and functional similarity/profile similarity (whole-brain averages: ρ_children_−ρ_adolescents_=.16, p=.06; ρ_children_−ρ_adolescents_ =.06, p=.06).

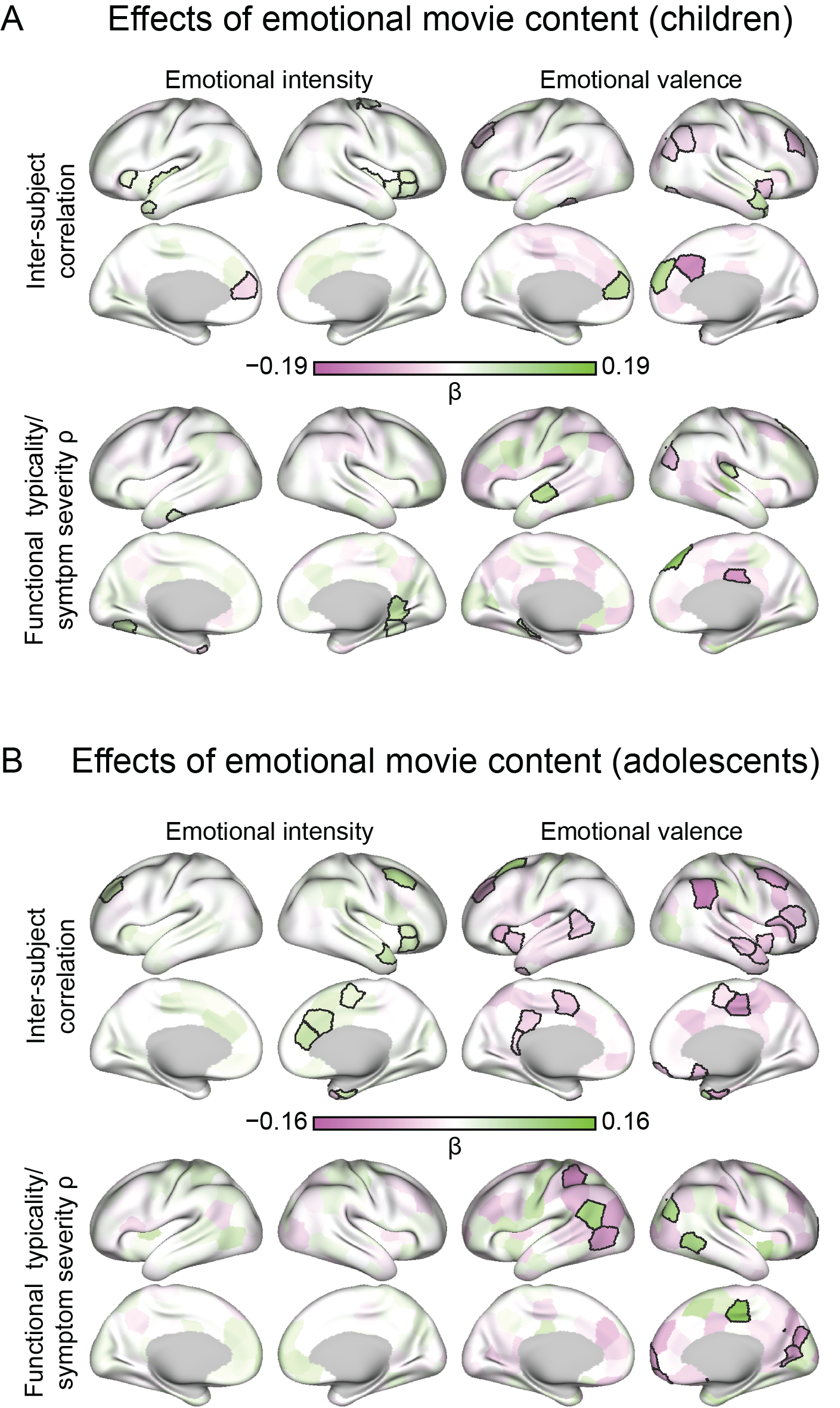

**Figure S2: Fluctuations in emotional movie content influence both inter-subject correlations and the functional typicality/depressive symptom severity relationship.** (A) In children, inter-subject correlations were greatest during more emotionally-charged moments of the movie when the (negative) typicality-depression score relationship was weakest (as determined by fGLS regression; bordered regions significant at p<.01, uncorrected). Emotional valence was differentially related to both increased BOLD synchronization and the functional typicality/symptom severity relationships. (B) The same analyses in (A) for the adolescent group show largely consistent effects. In adolescents, emotional valence was predominately negatively associated with ISC, while no relationships between emotional intensity and the strength of the functional typicality/symptom severity passed the p<.01 threshold.

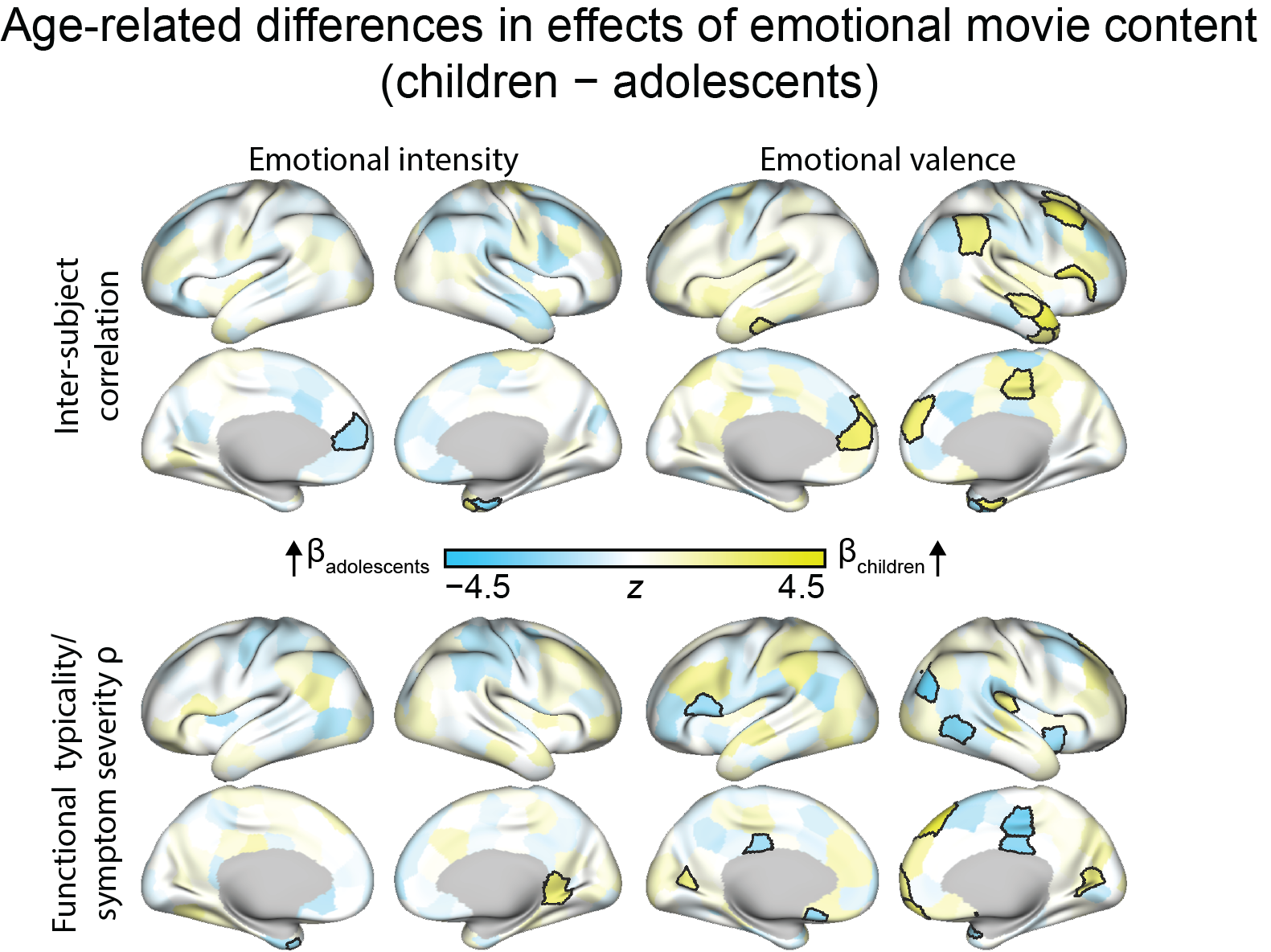

**Figure S3: Age-related differences (children – adolescents) in how emotional movie content influences inter-subject synchronization and the functional typicality/depressive symptom severity relationship.** These maps indicate the difference in fGLS-derived beta coefficients between the two age groups for each of the analyses shown in Figure S2 (p<.01, uncorrected).

**Tables S2-7: Full sample results (p<.01)**

**Table S2: Functional Typicality / Depressive Symptom Severity**

| parcel # | volume mm^3^ | MNI x | MNI y | MNI z | spearman’s ρ | p value |
| --- | --- | --- | --- | --- | --- | --- |
| 193 | 6128 | -60 | -27 | -18 | -.2829 | .0007 |
| 149 | 6700 | -39 | 17 | 47 | -.2706 | .0019 |
| 154 | 6106 | -43 | 42 | 11 | -.2550 | .0034 |
| 230 | 2470 | -32 | -40 | -4 | -.2434 | .0047 |
| 61 | 8377 | 59 | -3 | 3 | -.2417 | .0059 |
| 70 | 7893 | 61 | -43 | -18 | -.2336 | .0069 |
| 147 | 7059 | -46 | 28 | 27 | -.2307 | .0072 |
| 254 | 3363 | -21 | -53 | -24 | -.2282 | .0080 |
| 220 | 1376 | -4 | -5 | 33 | .2356 | .0083 |

**Table S3: Functional Similarity / Depressive Symptom Profile Similarity**

| parcel # | volume mm^3^ | MNI x | MNI y | MNI z | spearman’s ρ | p value |
| --- | --- | --- | --- | --- | --- | --- |
| 182 | 7464 | -42 | -66 | 42 | -.0831 | .0068 |
| 230 | 2470 | -32 | -40 | -4 | -.1137 | .0072 |

**Table S4: Inter-subject Synchronization / Emotional Intensity**

| parcel # | volume mm^3^ | MNI x | MNI y | MNI z | beta | p value |
| --- | --- | --- | --- | --- | --- | --- |
| 20 | 3744 | 37 | 21 | 6 | .1105 | .0001 |
| 14 | 5968 | 41 | 14 | 48 | .0899 | .0001 |
| 37 | 4568 | 38 | -12 | -1 | .0709 | .0003 |
| 24 | 4343 | 6 | -22 | 66 | .0788 | .0006 |
| 268 | 2439 | -6 | -19 | -37 | .0697 | .0009 |
| 169 | 3888 | -39 | 8 | -5 | .0563 | .0011 |
| 28 | 3970 | 6 | 14 | 49 | .0763 | .0011 |
| 36 | 4090 | 37 | 21 | -10 | .0810 | .0012 |
| 218 | 4170 | -8 | -22 | 46 | .0754 | .0012 |
| 155 | 4516 | -32 | 22 | 6 | .0630 | .0016 |
| 83 | 4695 | 8 | 35 | 17 | .0652 | .0019 |
| 9 | 8091 | 29 | 51 | 19 | .0616 | .0028 |
| 194 | 3027 | -49 | -5 | -37 | .0560 | .0034 |
| 15 | 5070 | 7 | 21 | 31 | .0679 | .0036 |
| 243 | 2544 | -19 | -46 | -53 | .0560 | .0050 |
| 186 | 3885 | -35 | 19 | -32 | .0533 | .0085 |
| 52 | 4094 | 40 | 19 | -34 | .0558 | .0089 |

**Table S5: Inter-subject Synchronization / Emotional Valence**

| parcel # | volume mm^3^ | MNI x | MNI y | MNI z | beta | p value |
| --- | --- | --- | --- | --- | --- | --- |
| 236 | 3147 | -7 | -66 | -38 | -.2033 | .0001 |
| 34 | 3555 | 42 | 5 | -8 | -.1111 | .0001 |
| 110 | 4146 | 21 | -55 | -24 | .1184 | .0001 |
| 122 | 2597 | 14 | -4 | 21 | .1078 | .0002 |
| 11 | 6389 | 38 | 35 | 31 | -.1181 | .0002 |
| 256 | 1749 | -24 | -38 | -44 | -.1086 | .0002 |
| 15 | 5070 | 7 | 21 | 31 | -.1046 | .0002 |
| 59 | 3171 | 43 | -26 | -25 | -.1066 | .0003 |
| 10 | 7549 | 8 | 53 | 24 | .1091 | .0004 |
| 123 | 3729 | 13 | 20 | -1 | .1047 | .0005 |
| 47 | 9524 | 54 | -45 | 37 | -.0928 | .0006 |
| 115 | 3194 | 8 | -57 | -51 | .0755 | .0010 |
| 155 | 4516 | -32 | 22 | 6 | -.0851 | .0019 |
| 24 | 4343 | 6 | -22 | 66 | -.0775 | .0022 |
| 201 | 4180 | -47 | -40 | -24 | -.0970 | .0027 |
| 32 | 3273 | 32 | -5 | 52 | .0861 | .0034 |
| 265 | 3632 | -5 | -22 | -16 | .0842 | .0046 |
| 173 | 3446 | -41 | -16 | 14 | -.0855 | .0067 |
| 268 | 2439 | -6 | -19 | -37 | .0908 | .0070 |
| 49 | 6082 | 41 | -75 | 28 | -.0776 | .0074 |
| 146 | 7833 | -27 | 34 | 36 | -.0650 | .0078 |
| 114 | 4651 | 23 | -72 | -29 | -.0755 | .0082 |
| 220 | 1376 | -4 | -5 | 33 | -.0905 | .0084 |
| 20 | 3744 | 37 | 21 | 6 | -.0864 | .0092 |

**Table S6: Functional Typicality & Symptom Severity Relationship / Emotional Intensity**

| parcel # | volume mm^3^ | MNI x | MNI y | MNI z | beta | p value |
| --- | --- | --- | --- | --- | --- | --- |
| 5 | 6436 | 8 | 46 | -2 | .0762 | .0005 |
| 249 | 3934 | -35 | -50 | -54 | .0895 | .0007 |
| 83 | 4695 | 8 | 35 | 17 | .0813 | .0014 |
| 138 | 4749 | -7 | 48 | -6 | .0939 | .0020 |
| 92 | 3590 | 31 | 4 | -22 | -.0650 | .0022 |
| 84 | 3038 | 5 | -1 | 36 | .0759 | .0030 |
| 86 | 4903 | 12 | -57 | 18 | .0613 | .0038 |
| 63 | 6510 | 62 | -24 | -3 | .0573 | .0045 |
| 55 | 5545 | 61 | -23 | -22 | .0668 | .0054 |
| 153 | 3883 | -32 | 20 | -16 | .0649 | .0077 |
| 207 | 6059 | -26 | -63 | -12 | .0714 | .0081 |
| 190 | 4258 | -58 | -6 | -23 | .0713 | .0084 |
| 9 | 8091 | 29 | 51 | 19 | .0604 | .0091 |
| 222 | 5645 | -9 | -59 | 18 | .0658 | .0093 |
| 141 | 5640 | -12 | 65 | 4 | .0624 | .0094 |

**Table S7: Functional Typicality & Symptom Severity Relationship / Emotional Valence**

| parcel # | volume mm^3^ | MNI x | MNI y | MNI z | beta | p value |
| --- | --- | --- | --- | --- | --- | --- |
| 166 | 3471 | -28 | -9 | 56 | -.1811 | .0001 |
| 101 | 5245 | 6 | -51 | -12 | -.1422 | .0001 |
| 179 | 6156 | -36 | -39 | 48 | -.1218 | .0001 |
| 237 | 2705 | -9 | -51 | -40 | .1316 | .0002 |
| 144 | 6283 | -29 | 50 | 22 | -.1039 | .0019 |
| 244 | 5232 | -7 | -50 | -11 | -.1210 | .0020 |
| 88 | 2103 | 7 | -19 | 30 | -.1124 | .0035 |
| 76 | 6226 | 18 | -83 | -11 | .0963 | .0042 |
| 198 | 3900 | -27 | -43 | -16 | .0911 | .0055 |
| 150 | 4360 | -5 | 18 | 46 | -.0940 | .0059 |
| 142 | 4098 | -29 | 54 | 3 | -.0974 | .0069 |
| 85 | 3200 | 5 | -39 | 27 | -.0847 | .0069 |
| 209 | 6172 | -48 | -67 | 1 | -.0849 | .0073 |
| 131 | 1608 | 6 | -22 | -42 | .1018 | .0084 |

**Tables S8-13: Child group results (p<.01)**

**Table S8: Functional Typicality / Depressive Symptom Severity**

| parcel # | volume mm^3^ | MNI x | MNI y | MNI z | spearman’s ρ | p value |
| --- | --- | --- | --- | --- | --- | --- |

(no results significant at p<.01)

**Table S9: Functional Similarity / Depressive Symptom Profile Similarity**

| parcel # | volume mm^3^ | MNI x | MNI y | MNI z | spearman’s ρ | p value |
| --- | --- | --- | --- | --- | --- | --- |

(no results significant at p<.01)

**Table S10: Inter-subject Synchronization / Emotional Intensity**

| parcel # | volume mm^3^ | MNI x | MNI y | MNI z | beta | p value |
| --- | --- | --- | --- | --- | --- | --- |
| 170 | 4595 | -38 | -13 | -1 | .0841 | .0001 |
| 243 | 2544 | -19 | -46 | -53 | .0726 | .0005 |
| 20 | 3744 | 37 | 21 | 6 | .0785 | .0009 |
| 37 | 4568 | 38 | -12 | -1 | .0504 | .0026 |
| 187 | 2833 | -49 | 11 | -31 | .0644 | .0039 |
| 34 | 3555 | 42 | 5 | -8 | .0513 | .0043 |
| 36 | 4090 | 37 | 21 | -10 | .0670 | .0044 |
| 140 | 5575 | -6 | 48 | 12 | -.0689 | .0052 |
| 155 | 4516 | -32 | 22 | 6 | .0567 | .0061 |
| 26 | 5891 | 26 | -13 | 66 | .0495 | .0100 |

**Table S11: Inter-subject Synchronization / Emotional Valence**

| parcel # | volume mm^3^ | MNI x | MNI y | MNI z | beta | p value |
| --- | --- | --- | --- | --- | --- | --- |
| 236 | 3147 | -7 | -66 | -38 | -.1889 | .0001 |
| 15 | 5070 | 7 | 21 | 31 | -.1754 | .0001 |
| 256 | 1749 | -24 | -38 | -44 | -.1345 | .0001 |
| 10 | 7549 | 8 | 53 | 24 | .1333 | .0001 |
| 34 | 3555 | 42 | 5 | -8 | -.0977 | .0001 |
| 11 | 6389 | 38 | 35 | 31 | -.1227 | .0001 |
| 140 | 5575 | -6 | 48 | 12 | .1259 | .0001 |
| 268 | 2439 | -6 | -19 | -37 | .1009 | .0003 |
| 119 | 3861 | 30 | -36 | -31 | .1013 | .0006 |
| 146 | 7833 | -27 | 34 | 36 | -.0918 | .0008 |
| 52 | 4094 | 40 | 19 | -34 | .0984 | .0015 |
| 53 | 6898 | 53 | 11 | -22 | .0981 | .0016 |
| 122 | 2597 | 14 | -4 | 21 | .0872 | .0016 |
| 123 | 3729 | 13 | 20 | -1 | .0637 | .0020 |
| 128 | 4408 | 5 | -10 | 5 | .0642 | .0023 |
| 115 | 3194 | 8 | -57 | -51 | .0483 | .0037 |
| 131 | 1608 | 6 | -22 | -42 | .0710 | .0064 |
| 67 | 4550 | 36 | -69 | -17 | -.0586 | .0069 |
| 49 | 6082 | 41 | -75 | 28 | -.0752 | .0081 |
| 48 | 7536 | 48 | -62 | 35 | -.0638 | .0084 |
| 254 | 3363 | -21 | -53 | -24 | .0805 | .0086 |
| 201 | 4180 | -47 | -40 | -24 | -.0815 | .0087 |

**Table S12: Functional Typicality & Symptom Severity Relationship / Emotional Intensity**

| parcel # | volume mm^3^ | MNI x | MNI y | MNI z | beta | p value |
| --- | --- | --- | --- | --- | --- | --- |
| 98 | 5112 | 15 | -46 | 3 | .1215 | .0003 |
| 112 | 4877 | 20 | -74 | -50 | .0808 | .0020 |
| 207 | 6059 | -26 | -63 | -12 | .0983 | .0031 |
| 189 | 2641 | -23 | 9 | -39 | -.0825 | .0042 |
| 196 | 2411 | -52 | -18 | -29 | .0613 | .0047 |
| 68 | 5033 | 25 | -45 | -12 | .0594 | .0056 |
| 249 | 3934 | -35 | -50 | -54 | .0637 | .0057 |

**Table S13: Functional Typicality & Symptom Severity Relationship / Emotional Valence**

| parcel # | volume mm^3^ | MNI x | MNI y | MNI z | beta | p value |
| --- | --- | --- | --- | --- | --- | --- |
| 12 | 8570 | 14 | 37 | 49 | .1948 | .0001 |
| 88 | 2103 | 7 | -19 | 30 | -.1630 | .0001 |
| 110 | 4146 | 21 | -55 | -24 | .1561 | .0001 |
| 62 | 4081 | 40 | -26 | 14 | .1101 | .0017 |
| 197 | 6106 | -57 | -15 | -7 | .1382 | .0029 |
| 49 | 6082 | 41 | -75 | 28 | -.1095 | .0035 |
| 233 | 3611 | -21 | -31 | -11 | .1069 | .0078 |

**Tables S14-19: Adolescent group results (p<.01)**

**Table S14: Functional Typicality / Depressive Symptom Severity**

| parcel # | volume mm^3^ | MNI x | MNI y | MNI z | spearman’s ρ | p value |
| --- | --- | --- | --- | --- | --- | --- |
| 159 | 5095 | -58 | -6 | 27 | -.4436 | .0015 |
| 125 | 5085 | 14 | 8 | -9 | -.3884 | .0032 |
| 104 | 1543 | 24 | -36 | -43 | -.3813 | .0035 |
| 95 | 4512 | 28 | -28 | -14 | -.3774 | .0041 |
| 97 | 3004 | 25 | -3 | -31 | -.3765 | .0046 |
| 157 | 6830 | -46 | 8 | 29 | -.3748 | .0051 |
| 122 | 2597 | 14 | -4 | 21 | -.3522 | .0060 |
| 230 | 2470 | -32 | -40 | -4 | -.3461 | .0066 |
| 61 | 8377 | 59 | -3 | 3 | -.3545 | .0068 |
| 229 | 2338 | -21 | -37 | 6 | -.3495 | .0071 |
| 153 | 3883 | -32 | 20 | -16 | -.3425 | .0079 |
| 260 | 2379 | -15 | -4 | 21 | -.3503 | .0080 |
| 103 | 3493 | 14 | -40 | -25 | .3453 | .0083 |
| 193 | 6128 | -60 | -27 | -18 | -.3427 | .0097 |

**Table S15: Functional Similarity / Depressive Symptom Profile Similarity**

| parcel # | volume mm^3^ | MNI x | MNI y | MNI z | spearman’s ρ | p value |
| --- | --- | --- | --- | --- | --- | --- |
| 104 | 1543 | 24 | -36 | -43 | -.1342 | .0019 |
| 92 | 3590 | 31 | 4 | -22 | -.0942 | .0033 |
| 1 | 4965 | 14 | 57 | -17 | -.1039 | .0036 |
| 95 | 4512 | 28 | -28 | -14 | -.1872 | .0040 |
| 90 | 5764 | 6 | -57 | 38 | -.1639 | .0042 |
| 128 | 4408 | 5 | -10 | 5 | -.1069 | .0043 |
| 182 | 7464 | -42 | -66 | 42 | -.1232 | .0048 |
| 169 | 3888 | -39 | 8 | -5 | -.0901 | .0055 |
| 230 | 2470 | -32 | -40 | -4 | -.1634 | .0064 |
| 51 | 3019 | 27 | 12 | -39 | -.0939 | .0065 |
| 162 | 7259 | -9 | 0 | 67 | -.1252 | .0088 |
| 34 | 3555 | 42 | 5 | -8 | -.0784 | .0095 |
| 137 | 4548 | -8 | 40 | -21 | -.0999 | .0095 |

**Table S16: Inter-subject Synchronization / Emotional Intensity**

| parcel # | volume mm3 | MNI x | MNI y | MNI z | beta | p value |
| --- | --- | --- | --- | --- | --- | --- |
| 14 | 5968 | 41 | 14 | 48 | .1075 | .0001 |
| 268 | 2439 | -6 | -19 | -37 | .0867 | .0001 |
| 146 | 7833 | -27 | 34 | 36 | .0836 | .0001 |
| 97 | 3004 | 25 | -3 | -31 | .0893 | .0001 |
| 51 | 3019 | 27 | 12 | -39 | -.0763 | .0002 |
| 83 | 4695 | 8 | 35 | 17 | .0869 | .0003 |
| 53 | 6898 | 53 | 11 | -22 | .0826 | .0004 |
| 247 | 5762 | -10 | -82 | -32 | .0706 | .0004 |
| 128 | 4408 | 5 | -10 | 5 | .0841 | .0025 |
| 15 | 5070 | 7 | 21 | 31 | .0665 | .0029 |
| 36 | 4090 | 37 | 21 | -10 | .0739 | .0030 |
| 131 | 1608 | 6 | -22 | -42 | .0777 | .0040 |
| 20 | 3744 | 37 | 21 | 6 | .0714 | .0051 |
| 25 | 3690 | 7 | -8 | 53 | .0486 | .0098 |

**Table S17: Inter-subject Synchronization / Emotional Valence**

| parcel # | volume mm^3^ | MNI x | MNI y | MNI z | beta | p value |
| --- | --- | --- | --- | --- | --- | --- |
| 47 | 9524 | 54 | -45 | 37 | -.1567 | .0001 |
| 14 | 5968 | 41 | 14 | 48 | -.1275 | .0001 |
| 146 | 7833 | -27 | 34 | 36 | -.1285 | .0001 |
| 164 | 7552 | -23 | 11 | 54 | .1292 | .0001 |
| 97 | 3004 | 25 | -3 | -31 | -.1240 | .0001 |
| 114 | 4651 | 23 | -72 | -29 | -.1027 | .0001 |
| 89 | 3531 | 8 | -23 | 45 | -.1333 | .0001 |
| 126 | 3726 | 10 | -27 | -2 | .0894 | .0001 |
| 265 | 3632 | -5 | -22 | -16 | .1116 | .0002 |
| 117 | 2783 | 7 | -54 | -34 | -.1208 | .0004 |
| 122 | 2597 | 14 | -4 | 21 | .1202 | .0004 |
| 155 | 4516 | -32 | 22 | 6 | -.1045 | .0004 |
| 169 | 3888 | -39 | 8 | -5 | -.0770 | .0004 |
| 60 | 3128 | 31 | 1 | -44 | .0907 | .0008 |
| 4 | 2385 | 16 | 34 | -23 | -.0909 | .0008 |
| 100 | 5424 | 32 | -78 | -40 | -.0951 | .0012 |
| 16 | 7575 | 54 | 25 | 1 | -.1065 | .0013 |
| 2 | 3367 | 10 | 18 | -19 | -.0899 | .0016 |
| 238 | 4468 | -37 | -53 | -31 | -.0855 | .0016 |
| 123 | 3729 | 13 | 20 | -1 | .0812 | .0020 |
| 124 | 4105 | 27 | 6 | 0 | -.0890 | .0026 |
| 34 | 3555 | 42 | 5 | -8 | -.0792 | .0027 |
| 51 | 3019 | 27 | 12 | -39 | .0913 | .0030 |
| 161 | 4319 | -6 | -4 | 48 | -.0766 | .0030 |
| 185 | 2681 | -38 | 6 | -38 | -.0870 | .0036 |
| 1 | 4965 | 14 | 57 | -17 | -.0900 | .0046 |
| 236 | 3147 | -7 | -66 | -38 | -.0709 | .0046 |
| 19 | 6344 | 48 | 36 | 15 | -.0843 | .0068 |
| 53 | 6898 | 53 | 11 | -22 | -.0828 | .0069 |
| 25 | 3690 | 7 | -8 | 53 | -.0466 | .0070 |
| 192 | 7466 | -58 | -47 | 5 | -.0715 | .0075 |
| 227 | 1691 | -7 | -42 | 13 | -.0507 | .0076 |
| 223 | 2879 | -5 | -36 | 32 | -.0666 | .0081 |
| 64 | 6013 | 56 | -9 | -14 | -.0789 | .0086 |

**Table S18: Functional Typicality & Symptom Severity Relationship / Emotional Intensity**

| parcel # | volume mm^3^ | MNI x | MNI y | MNI z | beta | p value |
| --- | --- | --- | --- | --- | --- | --- |

(no results significant at p<.01)

**Table S19: Functional Typicality & Symptom Severity Relationship / Emotional Valence**

| parcel # | volume mm^3^ | MNI x | MNI y | MNI z | beta | p value |
| --- | --- | --- | --- | --- | --- | --- |
| 89 | 3531 | 8 | -23 | 45 | .1611 | .0001 |
| 179 | 6156 | -36 | -39 | 48 | -.1523 | .0001 |
| 82 | 5351 | 15 | -68 | 8 | -.1251 | .0005 |
| 101 | 5245 | 6 | -51 | -12 | -.1516 | .0005 |
| 6 | 5627 | 15 | 65 | 4 | -.1283 | .0005 |
| 1 | 4965 | 14 | 57 | -17 | -.1195 | .0009 |
| 209 | 6172 | -48 | -67 | 1 | -.1276 | .0012 |
| 236 | 3147 | -7 | -66 | -38 | .1134 | .0016 |
| 77 | 4326 | 8 | -75 | 25 | -.1297 | .0027 |
| 183 | 7604 | -51 | -56 | 20 | .1218 | .0028 |
| 239 | 2492 | -9 | -55 | -52 | .1406 | .0030 |
| 69 | 5572 | 55 | -56 | -5 | .1183 | .0044 |
| 49 | 6082 | 41 | -75 | 28 | .1181 | .0057 |
| 102 | 7680 | 39 | -75 | -30 | -.1110 | .0082 |
| 263 | 4523 | -5 | -10 | 6 | .1014 | .0092 |
